# *HisCl1* regulates gustatory habituation in sweet taste neurons and mediates sugar ingestion in *Drosophila*

**DOI:** 10.1101/2024.05.06.592591

**Authors:** Haein Kim, Ziqing Zhong, Xinyue Cui, Hayeon Sung, Naman Agrawal, Tianxing Jiang, Monica Dus, Nilay Yapici

## Abstract

Similar to other animals, the fly, *Drosophila melanogaster*, reduces its responsiveness to tastants with repeated exposure, a phenomenon called gustatory habituation. Previous studies have focused on the circuit basis of gustatory habituation in the fly chemosensory system ^1,2^. However, gustatory neurons reduce their firing rate during repeated stimulation ^3^, suggesting that cell-autonomous mechanisms also contribute to habituation. Here, we used deep learning-based pose estimation and optogenetic stimulation to demonstrate that continuous activation of sweet taste neurons causes gustatory habituation in flies. We conducted a transgenic RNAi screen to identify genes involved in this process and found that knocking down *Histamine-gated chloride channel subunit 1* (*HisCl1)* in the sweet taste neurons significantly reduced gustatory habituation. Anatomical analysis showed that *HisCl1* is expressed in the sweet taste neurons of various chemosensory organs. Using single sensilla electrophysiology, we showed that sweet taste neurons reduced their firing rate with prolonged exposure to sucrose. Knocking down *HisCl1* in sweet taste neurons suppressed gustatory habituation by reducing the spike frequency adaptation observed in these neurons during high-concentration sucrose stimulation. Finally, we showed that flies lacking *HisCl1* in sweet taste neurons increased their consumption of high-concentration sucrose solution at their first meal bout compared to control flies. Together, our results demonstrate that HisCl1 tunes spike frequency adaptation in sweet taste neurons and contributes to gustatory habituation and food intake regulation in flies. Since HisCl1 is highly conserved across many dipteran and hymenopteran species, our findings open a new direction in studying insect gustatory habituation.

## RESULTS

### *HisCl1* regulates gustatory habituation in sweet taste neurons

The sense of taste allows animals to detect specific nutrients and avoid toxic compounds. The gustatory assessment of food is mainly regulated by the chemosensory neurons located in various taste organs ^4-6^. The fly, *Drosophila melanogaster*, can assess food quality via taste neurons located in the proboscis, legs, and wings ^7-13^. Stimulating sweet taste neurons in the labellum or legs triggers proboscis extension, followed by labellar opening and food ingestion ^14-17^. Taste neurons can alter their activity based on the metabolic state or when exposed continuously to a particular nutrient ^18,19^. For example, sweet taste neurons reduce their responsivity to sugars when flies are kept on a high-sugar diet ^20^. These neurons also reduce their firing rate during prolonged sugar stimulation (Figures S1A and S1B), leading to a decrease in the proboscis extension response, a phenomenon called gustatory habituation ^3^. Previous studies have focused on the circuit basis of gustatory habituation in flies ^2,18,19,21^. Here we investigated the cell-intrinsic factors that allow sweet taste neurons to adapt to prolonged sensory stimuli.

To study gustatory habituation, we stimulated sweet taste neurons using a red-shifted channelrhodopsin, Chrimson, in hungry flies walking on an air-suspended spherical treadmill (Figure 1A). In these experiments, we chose to use optogenetic activation instead of sugar stimulation to minimize the effects of satiation upon sugar ingestion on the fly’s gustatory responsiveness. To automatically track the movements of the proboscis, we used DeepLabCut™, a deep learning-based pose estimation software ^22^. We used the coordinates of the head, rostrum, and haustellum to calculate the angle of the rostrum (θ) and used this metric to quantify gustatory habituation (Figures 1B and 1C). During the continuous activation (total duration=60s, constant LED) of sweet taste neurons (*Gr64f>Chrimson*), flies extended their proboscis, but this response was quickly abolished within seconds (Figure 1D and Video S1). In contrast, when sweet taste neurons were transiently activated (total duration=60s, 0.1Hz pulsed LED), flies extended their proboscis consistently upon each LED stimulus onset without showing any signs of habituation (Figure 1E and Video S2). These results confirmed that continuous activation of sweet taste neurons indeed leads to gustatory habituation.

**Figure 1.**
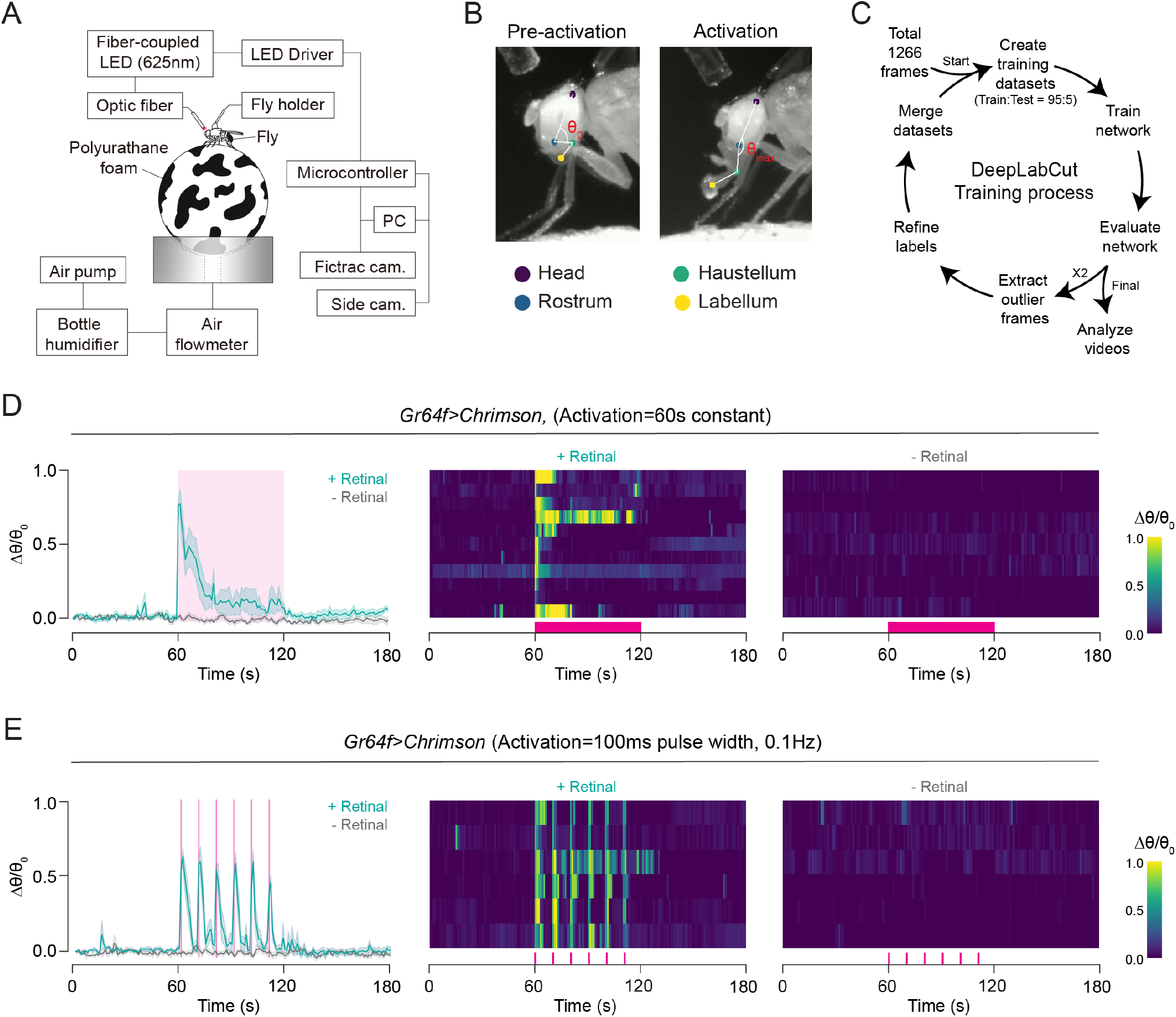
Continuous optogenetic activation of sweet taste neurons leads to gustatory habituation. (A) Schematic of the optogenetic stimulation and spherical treadmill setup. (B) Representative images for labeled body parts on the fly’s head (purple=Head, blue=Rostrum, green=Haustellum, and yellow=Labellum) and the rostrum angle (θ). (C) Schematic of the DeepLabCut™ pose estimation training process. (D-E) Change in rostrum angle (Δθ/θ_0_) is plotted over time (left, mean ± SEM) and as a heatmap (middle and right) in response to continuous (D) and pulsed (E) optogenetic activation (green=retinal fed flies, gray=no retinal controls, n=6-11). Magenta highlights represent when the LED light is ON for optogenetic activation. See also Videos S1 and S2.

To identify cell-intrinsic factors regulating gustatory habituation in flies, we conducted a transgenic RNAi screen by knocking down specific genes in sweet taste neurons during continuous optogenetic activation. In the mammalian brain, Ca^2+^-activated Cl^-^ channels (CACC) contribute to spike-frequency adaptation seen in the thalamocortical and hippocampal neurons ^23^. We hypothesized that a similar spike frequency adaptation mechanism might mediate gustatory habituation in the fly sweet taste neurons. To test our hypothesis, we first searched for chloride channel genes expressed in the fly proboscis using previously published RNA sequencing datasets ^20^. Next, using transgenic RNAi, we knocked down chloride channel genes (n=14) in the sweet taste neurons and tested these mutant flies in optogenetic activation experiments, identifying two genes, *HisCl1* and *Clc-a*, as candidate regulators of gustatory habituation (Figure 2A and Figure S1C). *Gr64f>HisCl1-RNAi* and *Gr64f>Clc-a-RNAi* flies consistently exhibited reduced gustatory habituation during optogenetic activation (Figures 1C and 2B). *Gr64f>HisCl1-RNAi* induced the most significant suppression in gustatory habituation; these flies continued to extend their proboscis despite continuous stimulation of sweet taste neurons (Figure 2C and Video S3). To confirm the RNAi knockdown results, we tested *HisCl1* null mutants (*HisCl1*^*-/-*^). When sweet taste neurons were continuously activated in *HisCl1*^*-/-*^ mutants, these flies showed a robust suppression in gustatory habituation similar to *Gr64f>HisCl1-RNAi* flies (Figures 1E and 2D). Interestingly, flies heterozygous for *HisCl1* (*HisCl1*^*+/-*^) behaved like hypomorphs; their gustatory habituation was not significantly different from *HisCl1*^*-/-*^ mutants or controls (Figure 2E).

**Figure 2.**
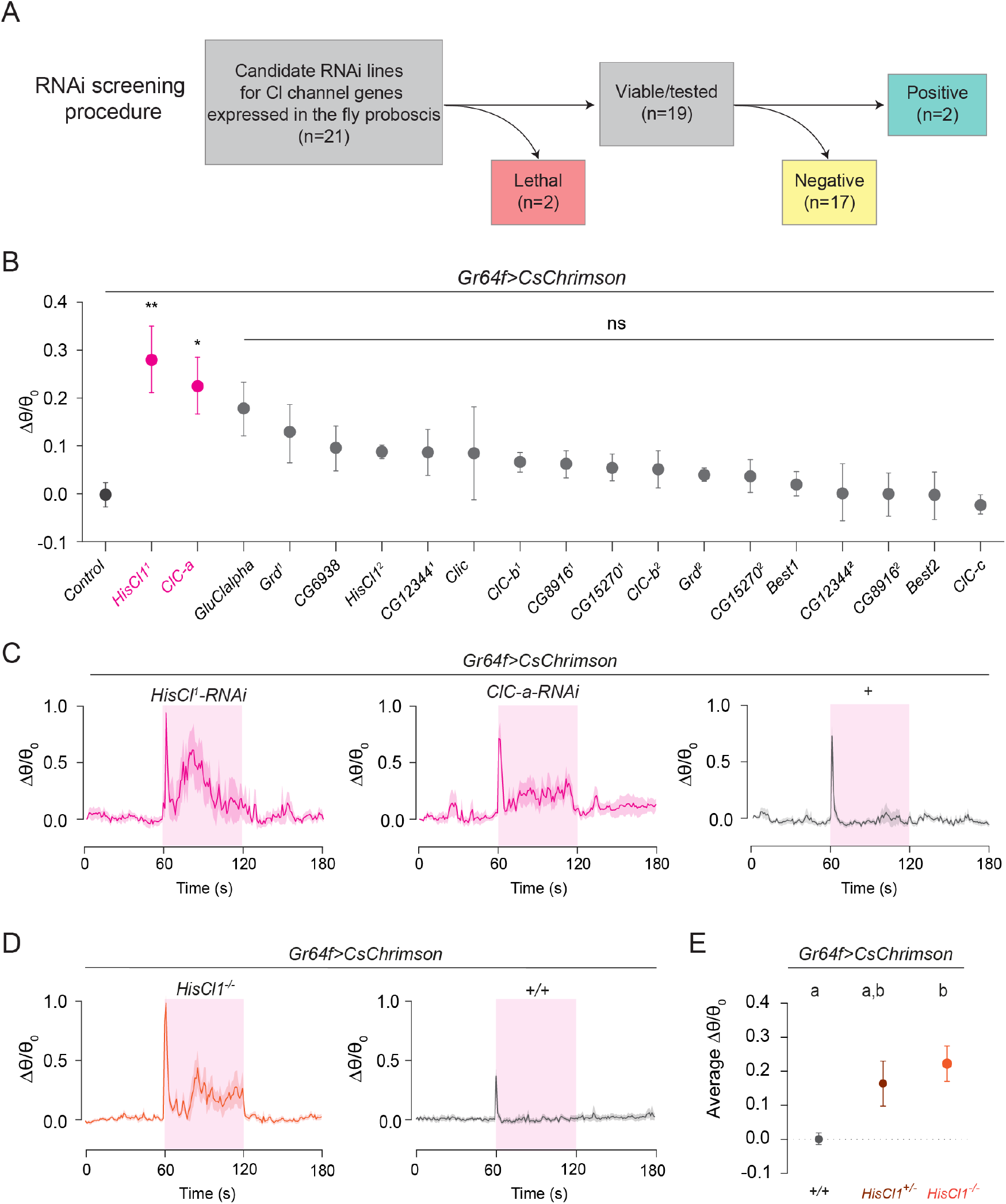
Knock-down of *HisCl1 and ClC-a* in sweet taste neurons suppresses gustatory habituation. (A) Outline of the transgenic RNAi screening procedure. (B) Average Δθ/θ_0_ for each RNAi line is quantified and plotted during continuous activation of sweet taste neurons. Magenta-labeled RNAi lines are significantly different from the control. (n=6-8, one-way ANOVA with Bonferroni test, *p<0.05, **p<0.01, ns = non-significant). (C-D) Δθ/θ_0_ is plotted over time (mean ± SEM, n=6-12) for indicated genotypes in response to continuous optogenetic activation. Magenta highlights represent when the LED light is ON for optogenetic activation. (E) Average Δθ/θ_0_ during optogenetic stimulation for *HisCl1*^*-/-*^ and controls in response to continuous activation of sweet taste neurons. (n=12, one-way ANOVA with Bonferroni test, genotypes labeled by different letters are statistically different from each other). See also Figure S1 and Video S3.

*HisCl1* gene encodes a histamine-gated chloride channel in flies. Two histamine-gated chloride channel genes are found in the *Drosophila* genome, *HisCl1* and *Ora transientless (Ort)* ^*24-26*^. HisCl1 and Ort work together in the fly visual system to mediate the inhibition between R7 and R8 photoreceptors ^27^. We hypothesized that Ort could also regulate gustatory habituation in sweet taste neurons similar to HisCl1. However, knocking down *Ort* did not impact gustatory habituation during the continuous activation of sweet taste neurons (Figures S2A and S2B). Consistent with our behavioral results, we found that GAL4 knock-in to the *Ort* gene *(Ort>*) did not label neurons in the fly chemosensory organs, while many optic lobe neurons were labeled in these flies (Figure S2C). Our results demonstrated that HisCl1, but not Ort, modulates gustatory habituation in sweet taste neurons.

### HisCl1 suppresses spike frequency adaptation in sweet taste neurons during high-concentration sugar stimulation

To further investigate how HisCl1 regulates gustatory habituation, we first examined its expression pattern in the fly taste organs. Using a GAL4 knock-in (*HisCl1>)*, we found that *HisCl1* is broadly expressed in the labellum (Figure 3A). Our detailed anatomical characterization showed that *HisCl1>* labeled a subpopulation of sweet taste neurons expressing the sugar receptor *Gr64f* ^17,28^. We observed 5.5 ± 0.5 neurons that are co-labeled by *HisCl1>* and *Gr64f>* in the labellum (n=6), 1.25 ± 0.48 in the labral sense organ (LSO) (n=4), and 0.5 ± 0.29 in the front tarsi (n=4) (Figure 3B). These results suggest that HisCl1+ and Gr64f+ cells contribute to gustatory habituation during continuous stimulation of sweet taste neurons.

**Figure 3.**
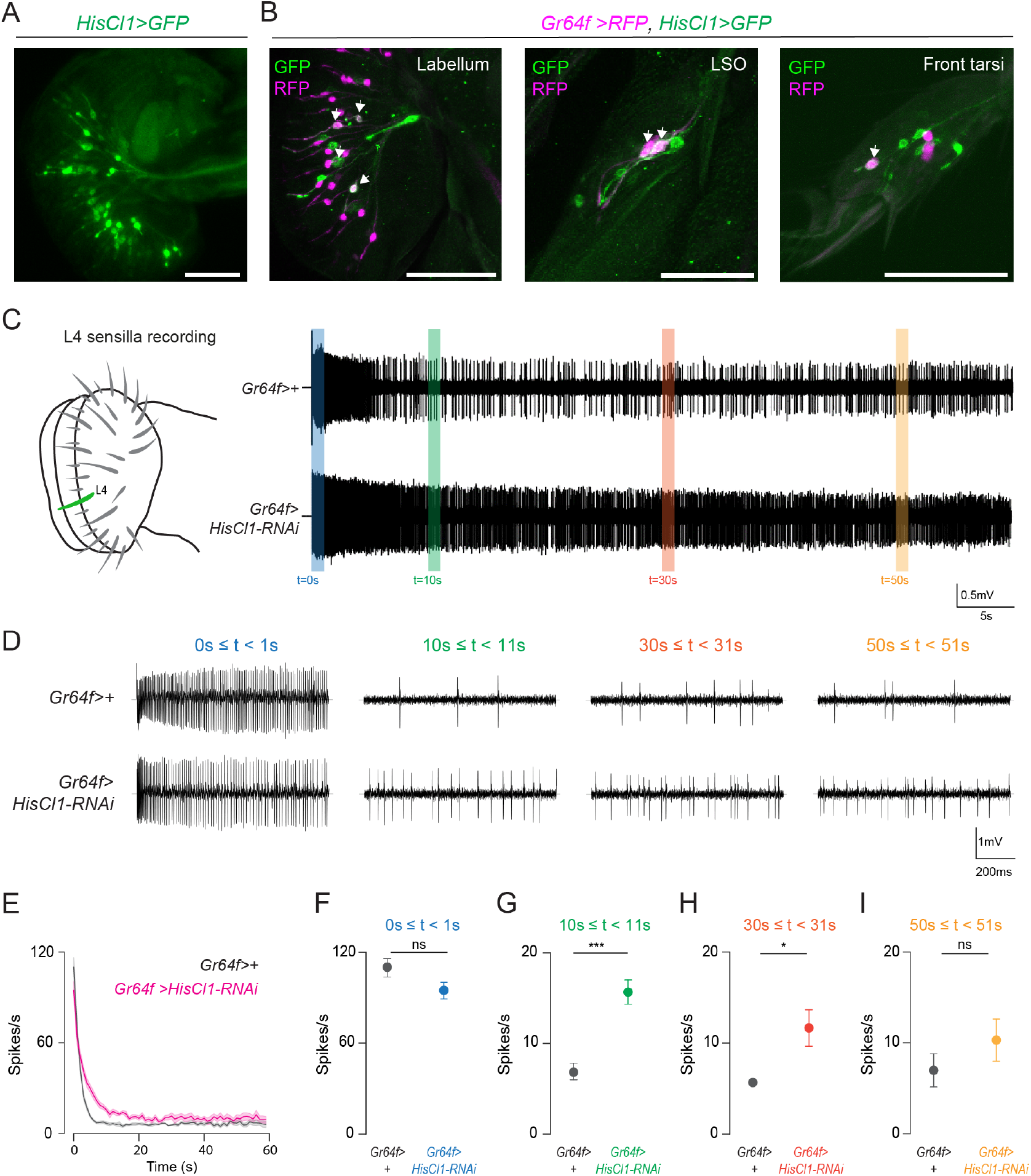
*HisCl1* is expressed in sweet taste neurons and regulates spike frequency adaptation during continuous sugar stimulation in these neurons. (A) Expression of *HisCl1>* (green) in the labellum (Scale bar=50µm). (B) Co-labeling of neurons expressing *Gr64f>* (magenta) and *HisCl1>* (green) in the labellum, LSO, and front tarsi. White arrows indicate Gr64+ and HisCl1+ neurons (Scale bars=50µm). (C) Representative single sensillum recording traces from L4 sensilla of control (top) and *Gr64f>HisCl1-RNAi* (bottom) flies during a 60-second 500mM sucrose stimulation. Colored vertical bars indicate time windows for firing rate quantifications in D. (D) Representative single sensillum recording traces from L4 sensilla of control (top) and *Gr64f>HisCl1-RNAi* (bottom) flies during 500mM sucrose stimulation in indicated time windows. (EB) Average L4 firing rates of control (gray) and *Gr64f>HisCl1-RNAi* (magenta) flies are plotted over time (mean ± SEM, n=6). (F-I) Average L4 firing rates of control (gray) and *Gr64f>HisCl1-RNAi* flies in indicated time windows (0-1s blue, 10-11s green, 30-31s orange, 50-51s yellow). Mean ± SEM, n=6, unpaired t-test with Welch’s correction, ns=non-significant, *p<0.05, ***p<0.001).

In flies, gustatory habituation is thought to be regulated by changes in the activity of taste circuits, mainly arising from postsynaptic partners of taste neurons^1,2^. However, sweet taste neurons also adapt their firing rate during continuous optogenetic stimulation ^3^ or during prolonged sugar exposure (Figures S1A and S1B), suggesting there should be changes in the intrinsic properties of these neurons leading to a depression in the firing rate. We hypothesize that HisCl1 might regulate gustatory habituation by directly altering sweet taste neuron firing rate during continuous stimulation. To directly test our hypothesis, we recorded the activity of HisCl1+ and Gr64f+ neurons located in the L4 sensillum in response to prolonged sucrose stimulation (Figure 3C). When the L4 sensillum was stimulated with 500mM sucrose, sweet taste neurons in both *Gr64f>HisCl1-RNAi* and control flies responded with a high initial firing rate that decreased within seconds, reaching a steady baseline (Figures 3C-3E). The firing rates reached a maximum between 0-1s after stimulus onset (Figures 3C and 3D). As we expected, the adaptation in firing rate was slower in *Gr64f>HisCl1-RNAi* flies compared to controls (Figures 3C and 3D, Figures S3C and 3D). To better quantify the temporal dynamics of neural activity during sugar stimulation, we compared the firing rates at early (0s≤t<1s), mid (10s≤t<11s and 30s≤t<31s), and late stages (50s≤t<51s) of the experiment (Figure 3D). The firing rates of sweet taste neurons in *Gr64f>HisCl1-RNAi* and control flies were not significantly different during the early or late stages of the sugar stimulation (Figures 3D, 3F, and 3I). However, during the mid-stages, while control flies rapidly adapted their firing rate after stimulus onset, *Gr64f>HisCl1-RNAi* flies continued to respond to the sugar stimulus with significantly higher firing rates (Figures 3D, 3G, and 3H). Interestingly, when we repeated the same experiment with 100mM sucrose solution, we found no differences in the firing rates of sweet taste neurons between the *Gr64f>HisCl1-RNAi* and control flies (Figures S3A and S3B). These results suggest that HisCl1 regulates spike frequency adaptation in sweet taste neurons, specifically during high-concentration sugar stimulation.

### HisCl1 is required in sweet taste neurons to regulate high-concentration sugar ingestion

Having established that HisCl1 regulates gustatory habituation and spike frequency adaptation in sweet taste neurons, we next explored how this behavioral and physiological adaptation impacts food intake behavior in flies. To quantify the temporal dynamics of food ingestion in *Gr64f>HisCl1-RNAi* and control flies, we used the Expresso automated food intake assay to capture meal-bouts of individual flies in real-time ^29^. We provided different concentrations of sucrose solution (20mM, 100mM, and 500mM) to 19-23hr food-deprived flies and recorded their ingestion behavior for 30 minutes (Figures 4A-4C). Our results showed that *Gr64f>HisCl1-RNAi* flies consumed significantly higher amounts of 500mM sucrose in their first meal bout compared to controls, while there were no differences in other concentrations tested between *Gr64f>HisCl1-RNAi* and control flies (Figures 4D-4F). We repeated the same experiment for *HisCl1*^*-/-*^ mutants and their genetic controls and found similar results (Figures S4A and S4B): *HisCl1*^*-/-*^ mutants consumed higher amounts of 500mM sucrose in their first meal bout compared to control flies (Figure S4C), and this difference in the meal bout volume was not observed for flies tested with 100mM sucrose (Figure S4C). In these experiments, we also quantified the total volume ingested for each genotype and found no significant differences among the groups tested (Figure 4G and Figure S4D). Our results demonstrate that HisCl1 function is required in sweet taste neurons to regulate meal bout volume when flies consume high concentration sucrose. Our findings from optogenetic activation, single sensillum recording, and food intake experiments led us to conclude that HisCl1 function is required in sweet taste neurons to suppress excessive sugar intake by fine-tuning spike frequency adaptation in these neurons.

**Figure 4.**
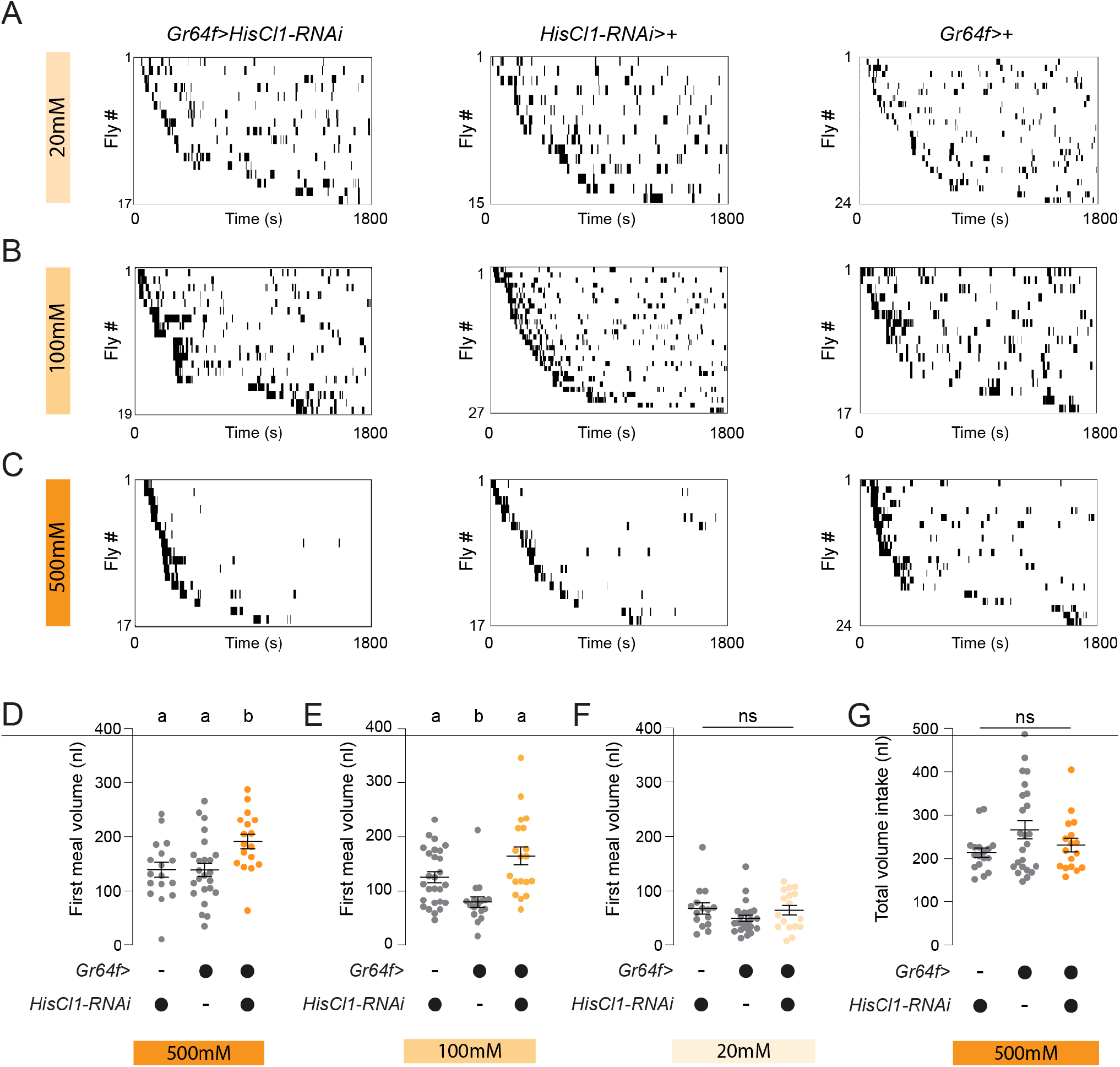
HisCl1 is required in sweet taste neurons to regulate sugar ingestion. (A-C) Meal bout raster plots of 19-23hr food-deprived flies of indicated genotypes ingesting 20mM (A), 100mM (B), or 500mM (C) sucrose solution in the Expresso. The trial duration is 30 minutes (n=15-27, indicated in the y-axis of each plot). (D-F) Average first meal bout volume of flies from indicated genotypes ingesting 500mM (D), 100mM (E), or 20mM (F) sucrose solution in the Expresso (n=15-27, mean ± SEM, one-way ANOVA with Bonferroni test. Groups labeled with different letters are significantly different, ns=non-significant). (G) The average total volume ingested for flies from indicated genotypes offered 500mM sucrose solution in the Expresso (n=17-24, mean ± SEM, one-way ANOVA, ns=non-significant).

## DISCUSSION

Gustatory habituation and dishabituation have been studied in *Drosophila* in the context of learning/memory and on the level of neural circuits ^1^. Here, we explored the cell-intrinsic mechanisms that regulate gustatory habituation in sweet taste neurons. We found a novel function for the histamine-gated chloride channel, HisCl1, in regulating temporal dynamics of sugar ingestion in hungry flies by fine-tuning the activity of sweet taste neurons. Interestingly, knocking down *HisCl1* in sweet taste neurons specifically suppressed gustatory habituation in response to high-concentration sucrose stimulation suggesting other cell-intrinsic factors also contribute to spike frequency adaptation in sweet taste neurons. Recently, it has been shown that a high-sugar diet decreases the stimulus-evoked firing rate and calcium responses of sweet taste neurons in flies due to the elevated activity of a conserved sugar sensor, O-linked N-Acetylglucosamine transferase (OGT) ^20^. It is possible that HisCl1 and OGT work together or in parallel pathways to fine-tune the activity of sweet taste neurons. Moreover, HisCl1 is expressed in a subpopulation of taste neurons, indicating there might be other cell-intrinsic factors regulating gustatory habituation. For example, in our genetic screen, we also identified another chloride channel, *Clc-a*, as a putative regulator of gustatory habituation. Our results suggest that distinct chloride channels might finetune the activity of different classes of taste neurons in flies.

How does HisCl1 regulate spike frequency adaptation in sweet taste neurons? In the mammalian brain, spike frequency adaption is regulated by Ca^2+^-activated chloride channels (CACC) ^23^. When neurons are strongly activated, an increase in the local Ca^2+^ levels activate CACC, leading to an influx of Cl^-^ and hyperpolarization of membrane potential, thereby decreasing the probability of spike generation ^30^. In the taste neurons of Necturus, an aquatic salamander, Ca^2+^-dependent Cl^-^ currents contribute to gustatory habituation ^31^. It is unclear if such Ca^2+^-dependent mechanisms trigger HisCl1 activation in the fly sweet taste neurons. Alternatively, HisCl1 might regulate gustatory habituation by changing synaptic release at the sweet taste neuron terminals. In the photoreceptors, HisCl1 plays a role in circadian entrainment via histamine signaling ^32,33^. Likewise, sugar exposure might trigger histamine release in downstream circuits, which activate HisCl1 in the sweet taste neuron axon terminals leading to presynaptic depression and spike frequency adaptation. Such phenomenon of self-backpropagation of presynaptic depression has been previously reported in hippocampal neurons^34,35^. To discriminate between these two possibilities, one needs to determine the subcellular localization of HisCl1 in sweet taste neurons and characterize its Cl-channel properties in response to the changes in intracellular Ca^2+^. Future experiments will address these open questions.

Our study also found that HisCl1 mutants increase their meal bout volume when ingesting a high-concentration sugar solution. We speculate that the increase in meal bout volume is mediated by the reduction in sensory adaptation in sweet taste neurons; when sweet taste neurons cannot reduce their firing rate during prolonged exposure to sugar, flies cannot cease ingestion leading to an increase in the meal bout volume. These findings suggest that one function of gustatory habituation in flies might be to avoid excessive sugar intake. Interestingly, HisCl1 is conserved across many dipteran and hymenopteran species, suggesting similar mechanisms might exist for regulating food ingestion in other insect species, including agricultural pests and human disease vectors. Overall, our study opens a new direction to studying the cell-intrinsic factors that regulate gustatory habituation in flies. We think this is a step forward in understanding how the insects adapt their behavior and sensory physiology when faced with persistent sensory stimuli.

## ACKNOWLEDGEMENTS

We thank the Yapici Lab members for their comments on the manuscript and Dr. David Anderson and Dr. Hubert Amrein for fly stocks. We acknowledge Bloomington Drosophila Stock Center (NIH P40OD018537) and the Developmental Studies Hybridoma Bank (NICHD of the NIH, University of Iowa) for reagents. Imaging data were acquired through the Cornell University Biotechnology Resource Center, with NIH S10OD018516 funding for the shared Zeiss LSM880 confocal/multiphoton microscope. Research in N.Y.’s lab is supported by a Pew Biomedical Scholar Award, the Alfred P. Sloan Foundation Award, and an NIH-R35-MIRA (R35GM133698). Research in M.D.’s lab is supported by a Klingenstein-Simons Fellowship in the Neurosciences, a Rita Allen Foundation Scholars award, and an NSF CAREER award (1941822).

## AUTHOR CONTRIBUTIONS

N.Y. and H.K. conceived the project and designed all the experiments. H.K. carried out and analyzed all the experiments except the electrophysiology recordings in Figure 3 and Expresso food intake quantification in Figure 4. E.Z. helped with experiments in Figure 1&2. N.A. helped with experiments in Figure S2. T. J. wrote the Python code for synchronizing the optogenetic stimulation with video recordings. N.Y., H.K, M.D., and H.S. designed the electrophysiology experiments in Figure 3 and Figure S4, which were then carried out and analyzed by H.S. and H.K. The Expresso food intake experiments in Figure 4 and Figure S3 were carried out and analyzed by X.C and H.K. N.Y. and H.K. interpreted the results and wrote the paper with feedback from other authors.

## DECLARATION OF INTERESTS

The authors declare no competing interests.

## SUPPLEMENTAL FIGURES, FIGURE TITLES, AND LEGENDS

**Figure S1.**
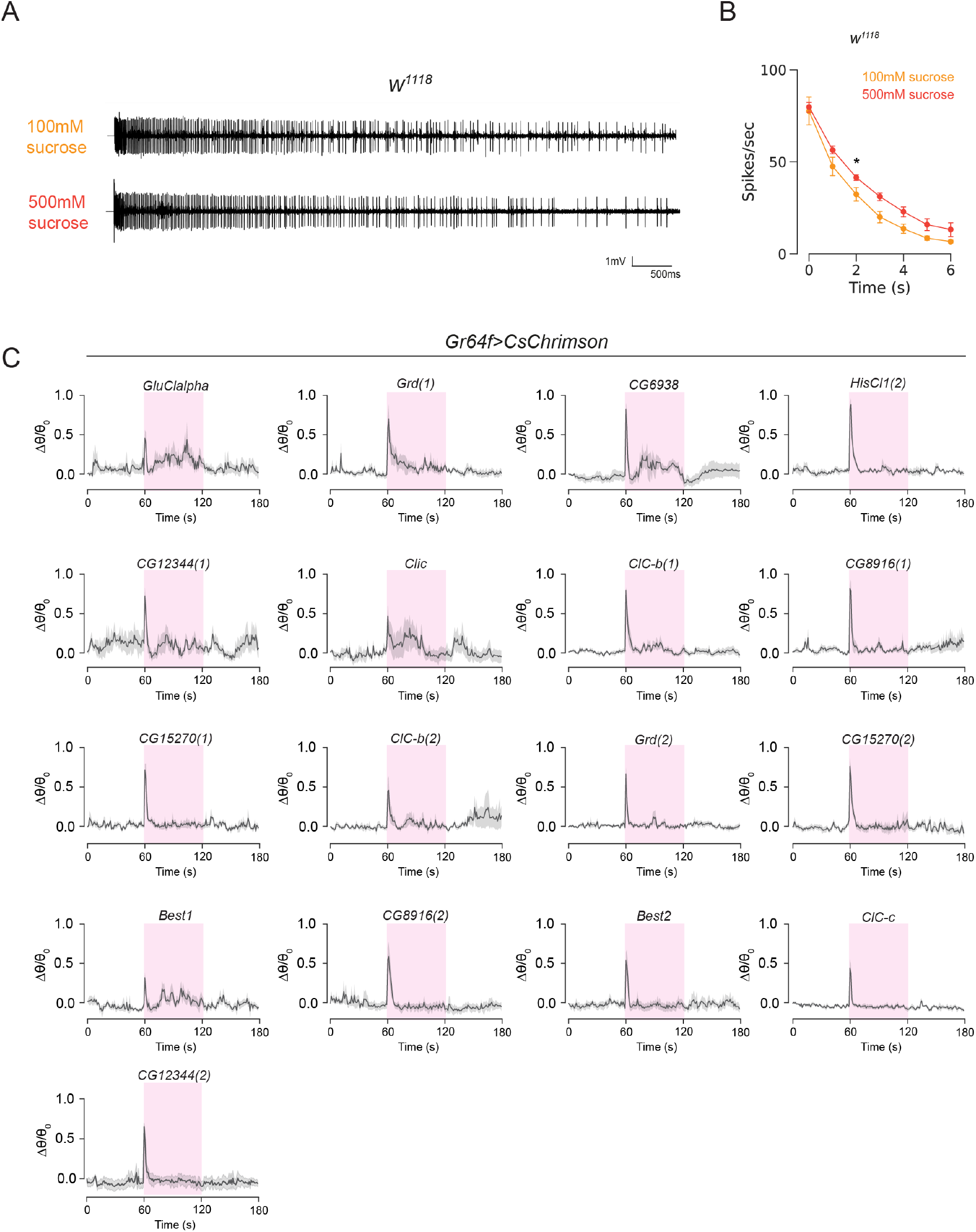
Single sensillum recording of taste neurons during continuous sugar stimulation and optogenetics data for negative RNAi lines targeting chloride channel genes. (A) Representative single sensillum recording traces from L4 sensilla of control flies during 100mM and 500mM sucrose stimulation. (B) The L4 firing rates during 100mM or 500mM sucrose stimulation are plotted over time (n=6-8, mean ± SEM, two-way ANOVA with Fisher’s pairwise comparison, * p<0.05). (C) Δθ/θ_0_ is plotted over time (mean ± SEM, n=6-8) for indicated genotypes in response to continuous optogenetic activation. Magenta highlights represent when the LED light is ON for optogenetic activation.

**Figure S2.**
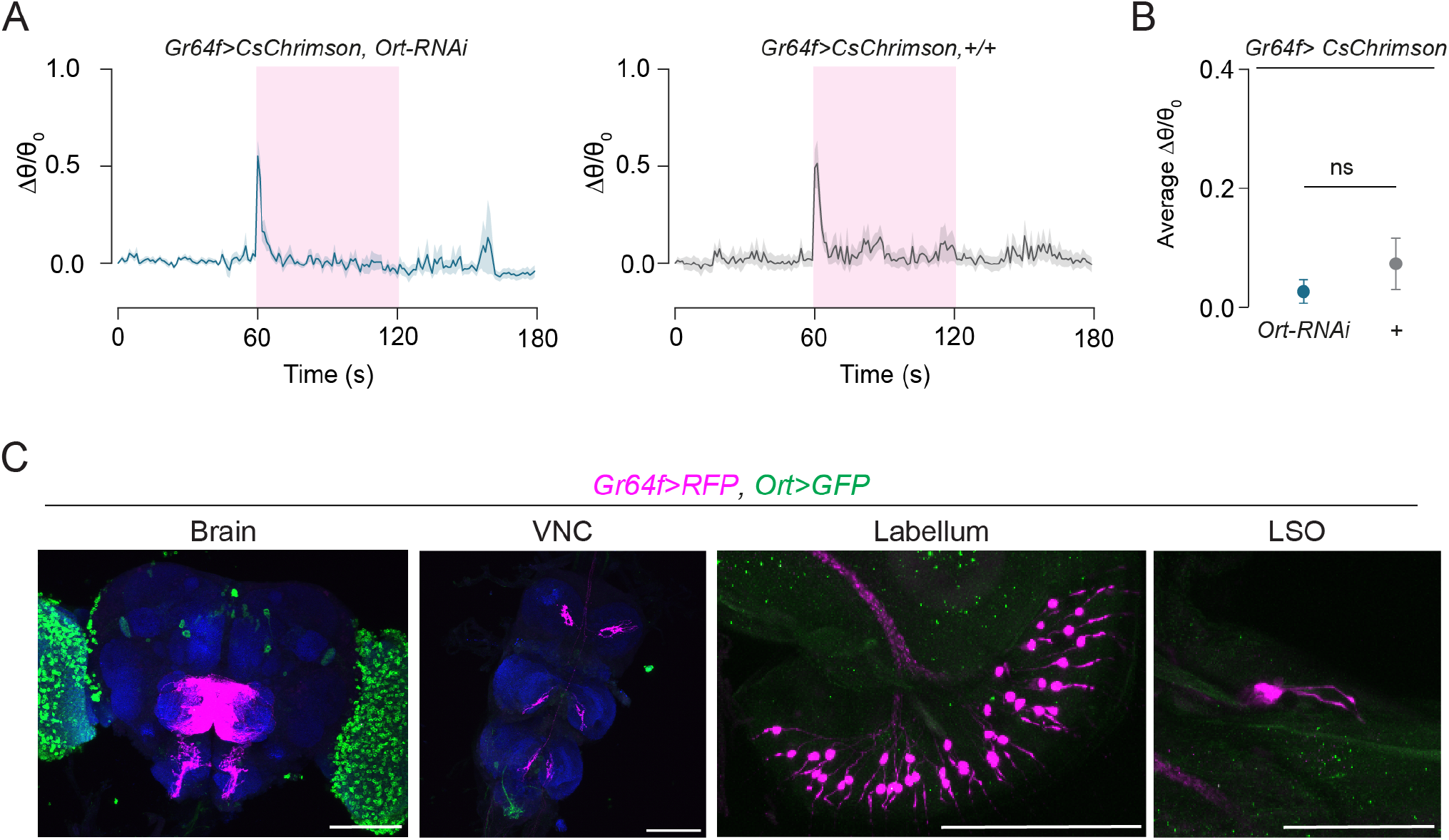
Knock-down of *Ort* in sweet taste neurons does not affect gustatory habituation. (A) Δθ/θ_0_ is plotted over time (mean ± SEM, n=6-8) for indicated genotypes in response to continuous optogenetic activation. Magenta highlights represent when the LED light is ON for optogenetic activation. (B) Average Δθ/θ_0_ during optogenetic stimulation for *Ort-RNAi* and control flies in response to continuous activation of sweet taste neurons (n=6-8, unpaired t-test with Welch’s corrections). (C) Labeling of neurons expressing *Gr64f>* (magenta) and *Ort>* (green) in the brain, VNC, labellum, and LSO. *Ort>* is expressed in the optic lobes and a few central neurons but not in any of the taste organs (Scale bars=50µm).

**Figure S3.**
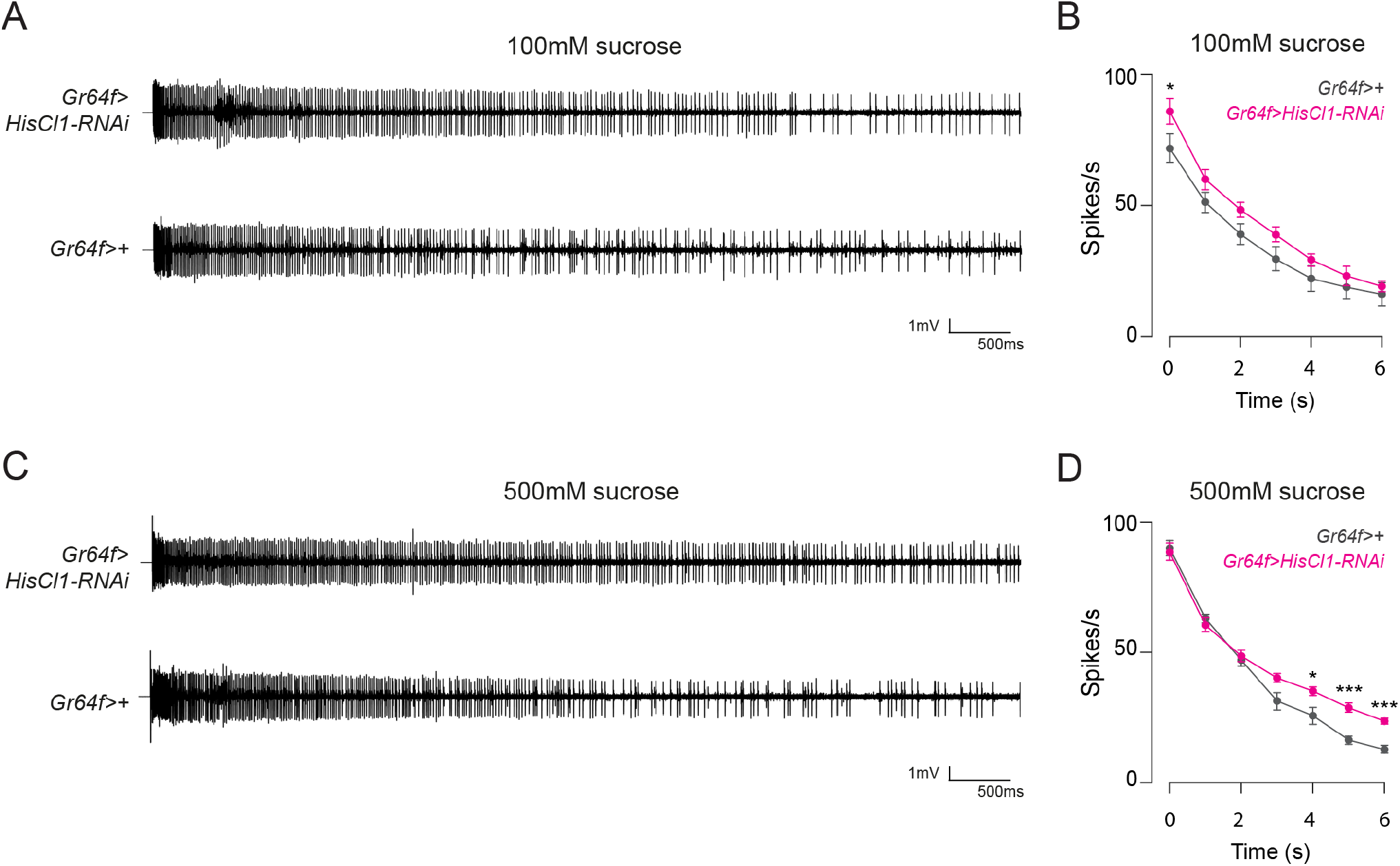
Knock-down of *HisCl1* in sweet taste neurons suppresses spike frequency adaptation during high-concentration sucrose stimulation. (A) Representative single sensillum recording traces from L4 sensilla of Gr64f>HisCl1-RNAi and control flies during 100mM sucrose stimulation. (B) >The L4 firing rates during 100mM sucrose stimulation are plotted over time for indicated genotypes (mean ± SEM, two-way ANOVA with Fisher’s pairwise comparison n=6-7). (C) Representative single sensillum recording traces from L4 sensilla of *Gr64f>HisCl1-RNAi* and control flies during 500mM sucrose stimulation. (D) The L4 firing rates during 500mM sucrose stimulation is plotted over time for indicated genotypes (mean ± SEM, two-way ANOVA with Fisher’s pairwise comparison n=6-7).

**Figure S4.**
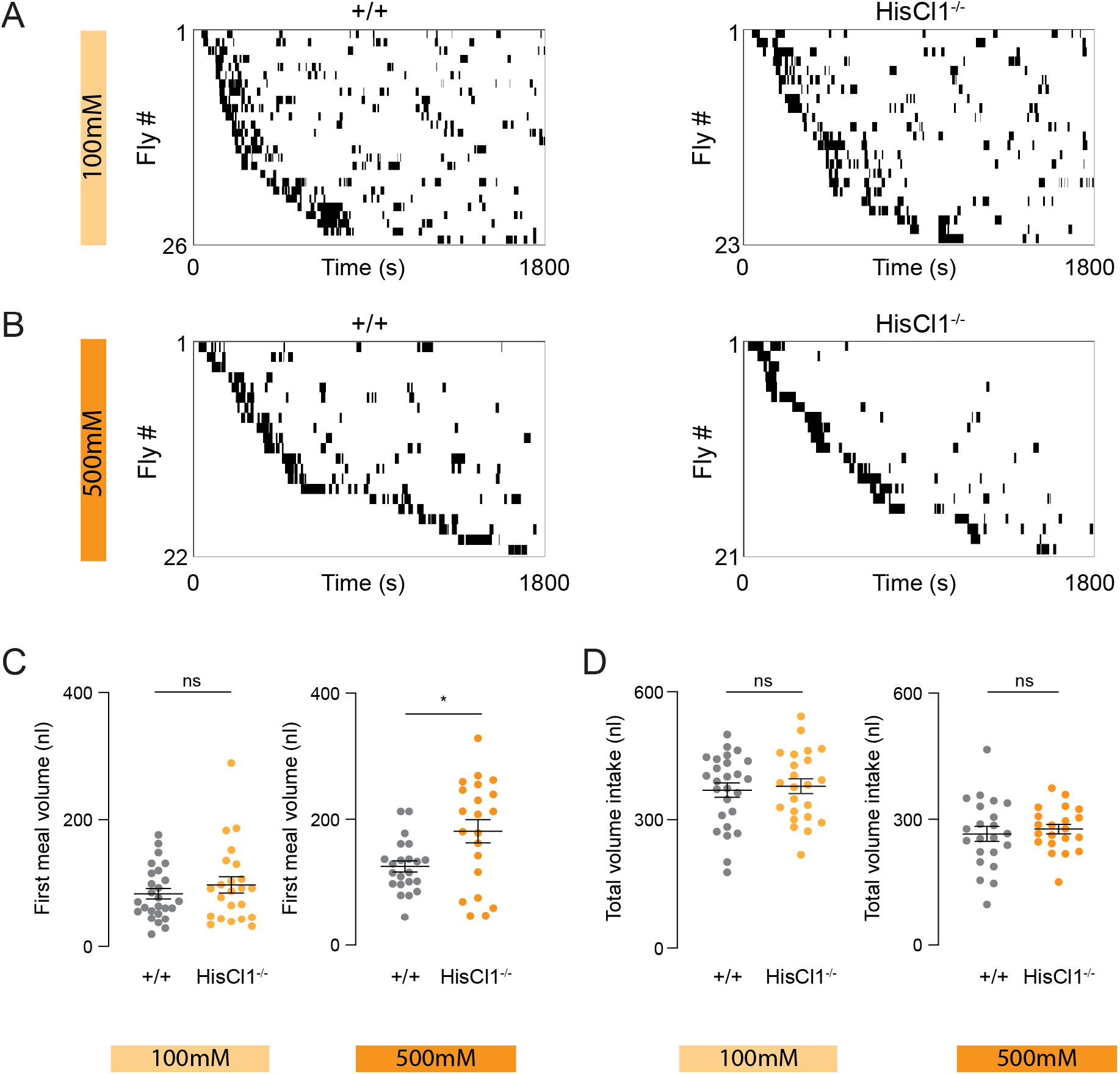
HisCl1 mutant flies increase their first bout volume while ingesting high-concentration sugar solution. (A-B) Meal bout raster plots of 19-23hr food deprived HisCl1 mutant or control flies ingesting 100mM (A) or 500mM (B) sucrose solution in the Expresso. (C) Average first meal bout volume of HisCl1 mutant or control flies ingesting 100mM, or 500mM sucrose solution in the Expresso (100mM n=23-26, 500mM n=21-22, mean ± SEM unpaired t-test with Welch’s correction, ns=non-significant, *p<0.05). (D) Average total meal bout volume ingested for HisCl1 mutant, or control flies offered 100mM, or 500mM sucrose solution in the Expresso (100mM n=23-26, 500mM n=21-22, mean ± SEM unpaired t-test with Welch’s correction, ns=non-significant, *p<0.05).

## STAR METHODS

## RESOURCE AVAILABILITY

### Lead contact

Further information and requests for resources and reagents should be directed to and will be fulfilled by the lead contact, Nilay Yapici.

### Materials availability

This study did not generate new unique reagents.

### Data and code availability

1. The data supporting this study’s findings are available from the lead contact upon reasonable request.
2. This study did not generate new code.
3. Any additional information required to reanalyze the data reported in this paper is available from the lead contact upon reasonable request.

## EXPERIMENTAL MODEL AND SUBJECT DETAILS

### Flies

*Drosophila melanogaster* was maintained on conventional cornmeal-agar-molasses medium at 25°C and 60-70% relative humidity under a 12hr light: 12hr dark cycle (lights on at 9 A.M.). Fly stocks and genotypes are detailed in Key Resources and Table S1. All fly stocks were obtained from Bloomington Drosophila Stock Center unless otherwise stated. The candidate genes targeted in the screen were selected for their functional annotation as Cl^-^ channels and their expression in the fly proboscis based on previous RNA sequencing experiments ^20^.

## METHOD DETAILS

### Optogenetic activation on the spherical treadmill

6–10-day old male flies were used in all optogenetic activation experiments. All flies were food deprived for 18-24 hours. During deprivation, test group flies were kept in a vial containing a Kimwipe (Kimtech Science™) soaked with 0.5mM all-trans-retinal (Sigma, R2500), and control flies were kept with a Kimwipe soaked with water. The spherical treadmill was custom-built from polyurethane foam (FR-7112, Last-A-Foam, General Plastics Manufacturing Company), as previously described ^36^. The ball had a 10mm diameter and a weight of 97mg, and it was air-floated by an air pump attached to a mass flow controller working at 0.45l/min. Two IR LED lights were used to illuminate the ball and the fly for better tracking quality (SZ-01-R8, Luxeon Star). The humidity of the air was maintained by passing it through a bottle humidifier (Salter Labs). During the optogenetic activation experiments, flies were tethered but allowed to walk on a spherical treadmill in the dark. Each trial started with a 60s recording of fly behavior without optogenetic stimulation, followed by 60s optogenetic stimulation, and another 60s recording without stimulation. The 625nm red LED light (Thorlabs, M625F2) was powered at 14µW/mm^2^ using a LED driver (Thorlabs, LEDD1B). During the optogenetic stimulation experiments, the red light was delivered to the fly using an optic fiber cannula (Thorlabs, CFMC22L20) (Figure 1). We used two optogenetic stimulation patterns; continuous activation (total duration=60s, LED constantly ON) and pulse activation (total duration=60s, LED 0.1Hz,100ms ON, 9900ms OFF). During the experiments, the movement of the fly’s proboscis was recorded using a Blackfly-S camera (BFS-U3-13Y3M-C, FLIR), and the fly/ball movements were recorded using a Firefly camera (FMVU-03MTM-CS, FLIR). All videos were acquired at 30fps. A custom-written script in Python controlled all video recordings and optogenetic stimulation patterns. We used FicTrac software ^37^ to extract the fly locomotion data by tracking the ball’s movements.

### Transgenic RNAi screen

Each transgenic RNAi line was crossed to flies carrying *Gr64f-GAL4* and *UAS-CsChrimson*-*mCherry*. To generate control flies, we crossed flies carrying *Gr64f-GAL4* and *UAS-CsChrimson*-*mCherry* to *w*^*1118*^. 6–10-day old male flies from the progeny were tested in the optogenetic activation experiments. All flies tested were food deprived for 24 hours in a vial containing a Kimwipe (Kimtech Science™) soaked with 0.5mM all-trans-retinal.

### Expresso Food Intake Quantification

Flies tested in the Expresso assay were prepared as described before ^29^. Briefly, 6–10-day old male flies were food deprived for 19-24 hours in a fly vial containing a piece of Kimwipe (Kimtech Science™) soaked with 1ml MilliQ water. On the day of the experiment, sucrose (Sigma, S5390) solutions were freshly prepared. Each fly was placed into a test cuvette and allowed to habituate for 5-10 mins before the start of the trial. Each trial lasted 30 mins, and the food intake data was recorded using the Expresso data acquisition software ^29^.

### Immunohistochemistry and Confocal Microscopy

Brain immunohistochemistry was carried out as previously described ^29^ with minor modifications. First, brains were dissected and fixed for 25 min with 4% paraformaldehyde (PFA) in PBST (PBS+0.3% Triton-X). After washing with PBST (4 times, 15 mins each), they were incubated with the blocking solution (5% normal goat serum (NGS, Jackson Labs, 005-000-121) in PBST) for 1.5 hours. Next, brains were incubated with the primary antibodies for two days at 4°C. After washing with PBST (3 times, 30 mins each), brains were incubated with the secondary antibodies for two days at 4°C. Finally, samples were mounted with SlowFade™ Gold Antifade Mountant (Thermo Fisher Scientific, S36936). Proboscis immunohistochemistry was carried out as previously described ^38^ with minor modifications. Proboscises were dissected and fixed with 4% PFA in PBST. After washing with PBST, samples were incubated with the blocking solution for 1.5 hours. Next, they were incubated with the primary antibodies for two days at 4°C. After washing with PBST (3 times, 30 min each), samples were incubated with the secondary antibodies for two days at 4°C. Samples were mounted with SlowFade™ Gold Antifade Mountant (Thermo Fisher Scientific, S36936). For imaging tarsi, legs were dissected and fixed with 4% PFA in PBST. After washing with PBST (3 times, 30 mins each), legs were mounted with SlowFade™ Gold Antifade Mountant (Thermo Fisher Scientific, S36936). Following antibodies were used for the immunohistochemistry: chicken anti-GFP (1:2000, Abcam, ab13970), rabbit anti-DsRed (1:500, Takara Bio, 632496), and mouse anti-Brp (1:20, DSHB, nc82), goat anti-chicken Alexa 488 (1:1000, Invitrogen, A-11039), goat anti-rabbit Alexa 546 (1:500, Invitrogen, A-11035), and goat anti-mouse Alexa 633 (1:500, Invitrogen, A-21052). All images were acquired using a Zeiss Confocal microscope (LSM 880) equipped with a 20X water immersion objective (Nikon, W Plan-Apochromat 20x/1.0). Confocal images were processed using the FIJI software.

### Single Sensillum Electrophysiology

We used the extracellular tip recording method to capture sucrose responses from the labellar taste sensilla as previously described ^39^. 6-10-day-old male flies were cold anesthetized, and the proboscis was immobilized by inserting the reference electrode containing the Beadle-Ephrussi Ringer solution through the thorax into the labellum. The neuronal firing rates of the L4 sensilla were recorded using a glass electrode (10-20 μm diameter) containing 100mM or 500mM sucrose mixed with an electrolyte (30 mM tricholine). The glass recording electrode was connected to the TastePROBE (Syntech) and the IDAC acquisition controller (Syntech). The signals were amplified (10x), band-pass-filtered (99-3000 Hz), and sampled at 12 kHz.

## QUANTIFICATION AND STATISTICAL ANALYSIS

### Proboscis Movement Quantification

The DeepLabCut software ^22^ was used to track the movements of the proboscis. The pose estimation model was trained by labeling the coordinates of four body parts on the fly’s head (head, rostrum, haustellum, and labellum) in each video frame. For the training process, we used 1266 frames to create training datasets. After the first round of training, the network was evaluated for errors by quantifying the pixel distance between the observed and estimated coordinates. The frames with errors were corrected and merged into the original training dataset, and the network was re-trained. After all the training, the final average error distance was less than 5 pixels. We used a custom-written script in Python for data analysis to calculate the rostrum angle (θ) using the three coordinates: head (x_hd,_ y_hd_), rostrum (x_r_, y_r_), and haustellum (x_h_, y_h_). Python script read all the output files from DeepLabCut as raw data, and the script collected three coordinates to compute the rostrum angle. Using three coordinates, two vectors were generated: 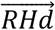 (head-rostrum) and 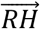 (rostrum-haustellum). To calculate the angle between the two vectors, we used the atan2 function that computes the counterclockwise angle between two vectors. Each rostrum angle was then averaged and aggregated in seconds. We calculated the Δθ/θ_0_ for all flies tested using the following formula, 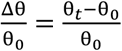 (θ_0_= initial angle t=0), θ_t_= angle at time (t), Δ θ= θ_t_ - θ_0_). Average Δθ/θ_0_ was calculated by averaging all values during the stimulation period (t=60-120s).

### Electrophysiology Data Analysis

To analyze neuronal firing rates in response to sugar stimulation, spikes were sorted manually, then the number of spikes per stimulation was determined using the Autospike software. Next, the total number of spikes was binned per second and plotted as a time series for each genotype. To quantify how neuronal firing rates changed between the early and the late phases of the recordings, average firing rates per genotype were calculated for the following time windows: short recordings (duration=6s); 0s ≤ t <3s and 4s ≤ t <7s), and long recordings (duration=60s); 0s ≤ t < 1s, 10s ≤ t < 11s, 30s ≤ t < 31s, and 50s ≤ t < 51s). The average spike rates in each time window were compared between the controls and *Gr64f>HisCl1* knock-down flies using the GraphPad Prism software.

### Expresso Food Intake Analysis

We used a custom-written code to analyze the Expresso food intake data ^40^. Each automatically detected meal bout was manually checked, and if there were mislabeled eating meal bouts in a trial, those flies were excluded from the data set. These mislabeled trials were less than ∼14% of total trials. We calculated the average total food intake and first meal bout volume of flies that have taken at least one meal bout during the assay period. Flies that did not consume food were not included in the quantifications. Data were plotted as scatter plots, indicating the mean ± SEM. Statistical analyses were performed using GraphPad Prism software.

## Notes

### Competing Interest Statement

The authors have declared no competing interest.

